# A novel hypomorphic allele of *Spag17* causes primary ciliary dyskinesia phenotypes in mice

**DOI:** 10.1101/2020.04.08.031393

**Authors:** Zakia Abdelhamed, Marshall Lukacs, Sandra Cindric, Heymut Omran, Rolf W. Stottmann

## Abstract

Primary ciliary dyskinesia (PCD) is a human condition of dysfunctional motile cilia characterized by recurrent lung infection, infertility, organ laterality defects, and partially penetrant hydrocephalus. We recovered a mouse mutant from a forward genetic screen that developed all the phenotypes of PCD. Whole exome sequencing identified this *primary ciliary dyskinesia only (Pcdo)* allele to be a nonsense mutation (c.5236A>T) in the *Spag17* coding sequence creating a premature stop codon at position 1746 (K1746*). The *Pcdo* variant abolished different isoforms of SPAG17 in the *Pcdo* mutant testis but not in the brain. Our data indicate differential requirements for SPAG17 in different motile cilia cell types. SPAG17 is required for proper development of the sperm flagellum, and is essential for either development or stability of the C1 microtubule structure within cilia, but not the brain ependymal cilia. We identified changes in ependymal cilia beating frequency but these did not apparently alter lateral ventricle cerebrospinal fluid (CSF) flow. Aqueductal (Aq) stenosis resulted in significantly slower and abnormally directed CSF flow and we suggest this is the root cause of the hydrocephalus. The *Spag17*^*Pcdo*^ homozygous mutant mice are generally viable to adulthood, but have a significantly shortened life span with chronic morbidity. Our data indicate that the c.5236A>T *Pcdo* variant is a hypomorphic allele of *Spag17* gene that causes phenotypes related to motile, but not primary, cilia. *Spag17*^*Pcdo*^ is a novel and useful model for elucidating the molecular mechanisms underlying development of PCD in the mouse.

## Introduction

Cilia are centriole-derived, microtubule-based membranous extensions that exist in almost every cell (Gilula and Satir 1972, Satir 2005, Pedersen, Veland et al. 2008). Ciliary structures are highly conserved across the animal kingdom. Largely based on the structure of the ciliary axoneme, cilia can be broadly classified into two main types. The motile cilia develop a central pair of microtubule singlets in the center of the axoneme and are therefore described to have a “9+2” arrangement of microtubule doublets (Nogales, Whittaker et al. 1999, Satir 2005). The primary cilia have a “9+0” structure of the axoneme (Nogales, Whittaker et al. 1999, Satir 2005) because they lack the central pair of microtubules and other molecular motors such as dynein arms and radial spokes which are responsible for ciliary movement. Therefore, primary cilia are nonmotile and mainly involved in mechanosensory functions (Pazour, Dickert et al. 2000, Goetz and Anderson 2010, Sánchez and Dynlacht 2016). The well documented function of the motile cilia is a coordinated rhythmic beating to move different body fluids in the brain, respiratory tract and the male and female genital ducts (Brightman and Palay 1963, Dirksen 1971, Jeffery and Reid 1975). This is consistent with the localized tissue distribution of motile cilia in these organs.

Defects in the assembly or function of motile cilia can cause primary ciliary dyskinesia (PCD). PCD (OMIM: 244400) is a rare and highly heterogeneous condition, with variants in over 40 causative genes reported to date (Bustamante-Marin, Yin et al. 2019, Cindrić, Dougherty et al., Lucas, Davis et al. 2019). In spite of this progress, a genetic diagnosis remains elusive in approximately 35% of PCD patients (Kurkowiak, Ziętkiewicz et al. 2015, Yang, Banerjee et al. 2018). PCD manifests as chronic respiratory tract infections, infertility, and laterality defects in about 50% of cases (Afzelius 1976, Afzelius 1981). These laterality defects develop due to defective nodal cilia in the developing embryo. Nodal cilia are a class of the motile cilia that lack the central pair apparatus but are equipped with inner and outer dynein arms. These enable the nodal cilia to produce a vortex-like movement to generate a gradient that establishes left–right asymmetry in the developing embryo (Nonaka, Tanaka et al. 1998, Marszalek, Ruiz-Lozano et al. 1999, Brueckner 2001). Hydrocephalus infrequently occurs in individuals with PCD and may reflect dysfunctional ependymal cilia, although various mechanisms have been proposed (Wessels, den Hollander et al. 2003, Wessels, den Hollander et al. 2003, Kosaki, Ikeda et al. 2004, Del Bigio 2010, Berlucchi, de Santi et al. 2012, Vieira, Lopes et al. 2012, Lee 2013). While there is not a clear link between nodal and ependymal cilia dysfunction, many reports indicate that hydrocephalus is over-represented in PCD patients as compared to the unaffected population. Intriguingly, almost all rodent models of PCD genes consistently develop hydrocephalus (Brody, Yan et al. 2000, Ibañez-Tallon, Pagenstecher et al. 2004, Lechtreck, Delmotte et al. 2008, Lee, Campagna et al. 2008, Lee, Campagna et al. 2008, Fernandez-Gonzalez, Kourembanas et al. 2009, Sironen, Kotaja et al. 2011, Ha, Lindsay et al. 2016, Abdelhamed, Vuong et al. 2018). Studies of rodent models of PCD on multiple inbred strains have begun to indicate that genetic modifiers influence susceptibility to PCD-associated hydrocephalus in mouse (Lee, Campagna et al. 2008, Sironen, Kotaja et al. 2011).

Ciliary motility is an ATP-dependent process that results in activation of dynein motor proteins between the pairs of microtubule doublets that are the major structural component of the ciliary axoneme (P Satir and Sleigh 1990). A significant number of studies indicate that signals from the central pair propagate through radial spokes to modulate the dynein activity and ultimately affect ciliary beating and flagellar movement (Adams, Huang et al. 1981, Smith and Yang 2004, Wirschell, Nicastro et al. 2009, Yang and Smith 2009, Zhu, Poghosyan et al. 2019). These conclude that dynein is a downstream effector of the central pair apparatus (Huang, Ramanis et al. 1982, Porter, Power et al. 1992, Porter, Knott et al. 1994, Rupp, Toole et al. 1996). The central pair consists of the two 13-protofilament microtubule singlets (C1 and C2) interconnected by bridge-like structure as well as several projections docked onto the C1 (C1a-C1f) and C2 (C2a-C2b) singlets. At least 23 polypeptides ranging in molecular weight from 14–360KDa compromise the central apparatus (Adams, Huang et al. 1981, Zhao, Hou et al. 2019). The C1 and C2 microtubule singlets and their associated projections are structurally and biochemically distinct. At least 10 different polypeptides are uniquely associated with the C1 microtubule, and seven are unique to the C2 microtubule (Dutcher, Huang et al. 1984, Lee 2011). This biochemical and structural asymmetry is believed to have functional consequences for cilia or flagellar beat and waveform (Inaba 2011). The 240KDa protein PF6 is the largest protein in the C1a projection (Rupp, O’Toole et al. 2001, Goduti and Smith 2012). Studies in *Chlamydomonas reinhardtii* indicate that PF6 acts as a scaffold protein essential for assembly of the smaller C1a components (Rupp, O’Toole et al. 2001), interacts with calcium binding protein calmodulin (Wargo, Dymek et al. 2005), and may play role in modulating the activity of both inner and outer dynein arms (Goduti and Smith 2012). As expected, *Chlamydomonas pf6* mutants develop paralyzed flagella and lack the central pair C1a projection (Rupp, O’Toole et al. 2001)

*Spag17* is the mammalian homologue of the *Chlamydomonas pf6* (Zhang, Jones et al. 2005). It is well documented the SPAG17 protein is present in motile cilia and flagella with a “9+2” axoneme structure (Zhang, Jones et al. 2005, Zhang, Zariwala et al. 2007) including the ciliated ependymal cells (Gokce, Stanley et al. 2016). Similar to its orthologue, the SPAG17 protein is present in the central pair apparatus of motile cilia and is essential for development of the C1a projection of the C1 microtubule singlet (Rupp, O’Toole et al. 2001, Zhang, Zariwala et al. 2007, Teves, Zhang et al. 2013). SPAG17 and other C1 proteins including SPAG6 and SPAG16L are known to bind within the C1a projection (Zhang, Jones et al. 2005, Zhang, Zariwala et al. 2007). *Spag17* null mice develop severe ciliary motility defects that lead to hydrocephalus, severe respiratory distress, and death within 12 hours after birth (Teves, Zhang et al. 2013). All of these phenotypes are indicative of motile cilia related abnormalities. Recently, *SPAG17* variants were reported to cause primary ciliary dyskinesia in human patients (Andjelkovic, Minic et al. 2018). These variants are linked to male infertility due to severe asthenozoospermia(Xu, Sha et al. 2018), and affect almost all stages of spermatogenesis in mice (Kazarian, Son et al. 2018).

The mammalian *Spag17* gene shows greater complexity in expression patterns and functions (Teves, Nagarkatti-Gude et al. 2016) than its orthologue *pf6*. Phenotypes beyond the motile cilia were also reported in the *Spag17* mutant mice which develop skeletal defects with shorter and less frequent primary cilia on the *Spag17* mutant chondrocytes, osteoblasts, and embryonic fibroblasts (MEFs; Teves, Sundaresan et al. 2015). This is consistent with genome wide association studies suggested that *SPAG17* may have a role in controlling human height trait (Weedon and Frayling 2008, Weedon, Lango et al. 2008, Takeuchi, Nabika et al. 2009, Kim, Lee et al. 2010, Zhao, Li et al. 2010, N’Diaye, Chen et al. 2011, van der Valk, for the Early Growth Genetics et al. 2014, Wood, Esko et al. 2014).

This study presents a careful phenotypic analysis of the novel *Spag17* hypomorphic allele we recovered from ENU induced forward genetic screen. We named this allele *primary ciliary dyskinesia only (Pcdo).* Our data indicate this *Spag17*^*Pcdo*^ allele is compatible with life but homozygous mutants have a shorter life span, and develop PCD phenotypes related due to defects in motile cilia function. We see no effects of the *Spag17*^*Pcdo*^ variant on primary cilia development or function in this model.

## Results

### *Pcdo* is a nonsense allele of *Spag17*

We identified the *primary ciliary dyskinesia only* (*Pcdo*) mutant in a mouse ENU mutagenesis forward genetic screen for recessive alleles leading to organogenesis phenotypes. ENU mutagenesis was performed as described previously (Stottmann, Moran et al. 2011, Stottmann and Beier 2014, Menke, Cionni et al. 2015, Lukacs, Roberts et al. 2019) and phenotyping to recover mutant alleles was performed at early postnatal stages as part of an experiment to look for mutants with abnormal forebrain development. *Pcdo* mutants were initially identified by enlarged head and distended lateral ventricles upon gross dissection. Further histological examination confirmed the occurrence of hydrocephalus and abnormal accumulation of mucus in the respiratory passages. There was no evidence of heterotaxy or any other abnormal organ laterality.

In order to identify the causal variant in *Pcdo* mutants, we performed whole exome sequencing on three phenotypic mutants, (Supplementary table 1). We filtered for variants which were homozygous for the alternate allele in each mutant, not present in dbSNP to rule out strain polymorphisms, predicted to have “high” or “moderate” impact on coding, and then common to all three sequenced mutants. Only three genes of predicted “high” impact met these criteria: *plexin C1* (*Plxnc1*), *Sfi1 homolog, spindle assembly associated (yeast)* (*Sfi1*) and *sperm associated antigen 17* (*Spag17*). A null allele of *Plxnc1* exists and does not have the same phenotype as the *Pcdo* mutants (Jeroen Pasterkamp, Peschon et al. 2003). *Sfi1* seems to be highly polymorphic as this routinely appears in similar analyses in our lab so we initially excluded this as a candidate. The null *Spag17* phenotype however, was sufficiently similar to *Pcdo* we initially pursued this as a candidate (Teves, Zhang et al. 2013)

*Spag17-204* (ENSMUST00000164539) consists of 49 exons and encodes for the full length 2320 amino acid sSPAG17 protein (ENSMUSP00000134066; Figure 1A). Another smaller splice variant encoding a 97KDa SPAG17 isoform is found in the testis. This is known to be proteolytically processed during the process of spermatogenesis and sperm maturation to generate 72KDa and 28KDa SPAG17 fragments (Zhang, Jones et al. 2005, Silina, Zayakin et al. 2011). The mouse SPAG17 protein has a small coiled coil domain followed by 2 regions of compositional bias with lysine-rich and Glycine-rich areas and C-terminal Pfam (flagellar associated PapD like) domains. The latter Pfam domain is highly conserved in mammals (Zhang, Jones et al. 2005; Figure 1A).

**Figure 1:**
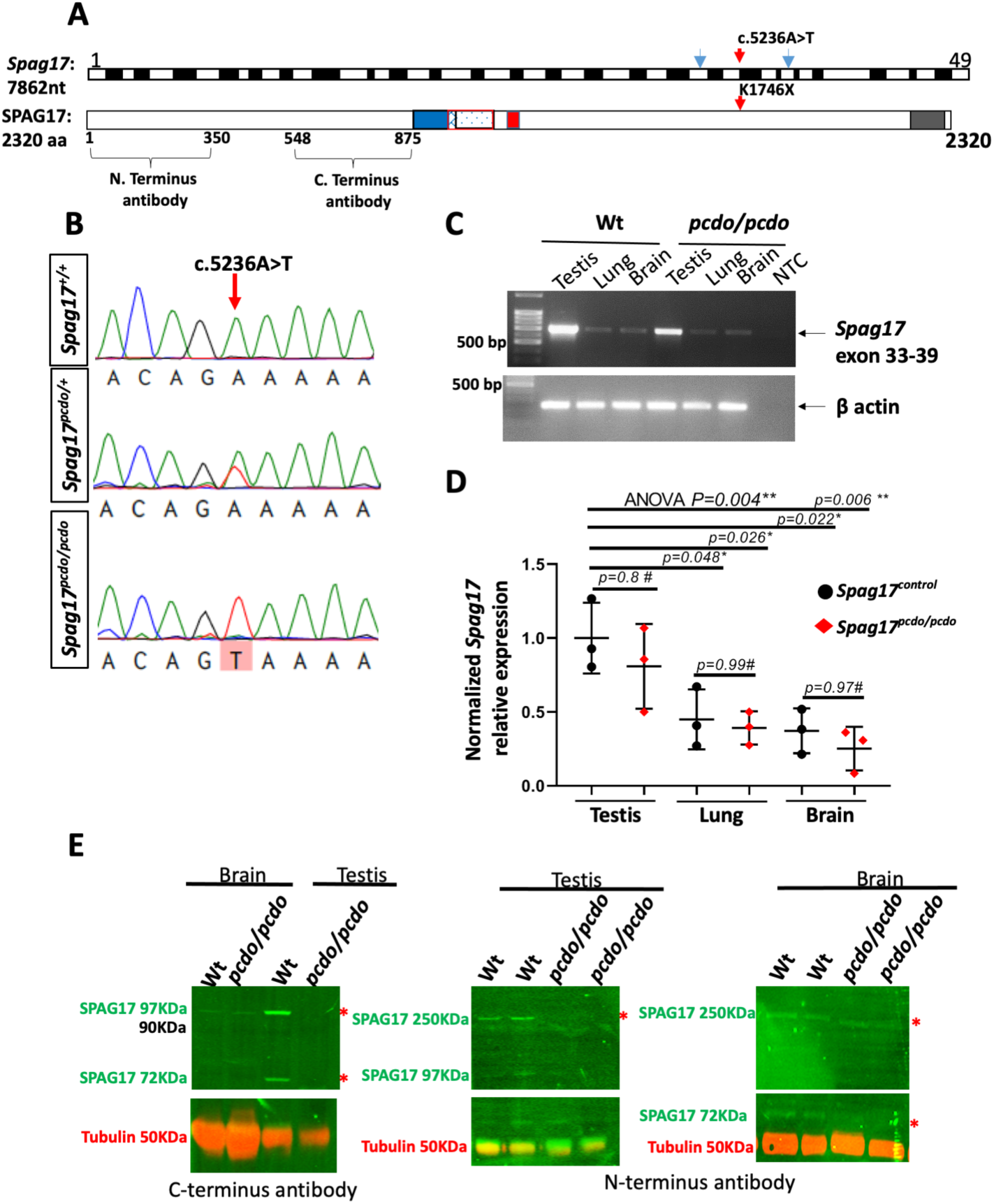
*Pcdo* is a nonsense allele of *Spag17*. The *Spag17* gene has 49 exons and position of the c.5236A>T in exon 36 is indicated with red arrow. Mouse SPAG17 is formed of 2320 amino acids, antigenic sites for the N. terminus and C. Terminus SPAG17 antibodies are indicated, blue arrows indicate positions of the primers used to assay expression of Spag17 shown in C (A). The mouse SPAG17 has coiled coil (blue), lysine-rich (blue dotts), Glycine-rich (red), and Pfam (grey)domains,the approximate position of the K1746* indicated with a red arrow (A). Electrophenographs of Sanger sequencing tracing showing the allele change A>T in the *Spag17*^*Pcdo/Pcdo*^ mutants (B). Semi-quantitative RT-PCR of the *Spag17* exon 33-39 and β. Actin loading control (C). Quantification of Spag17 expression relative to β. Actin loading in the testis, lung, and brain tissues, n=3 animals each genotype (D). Western blotting with anti-C. terminus (right), and N. terminus (middle, and left) SPAG17 antibodies on testis and brain lysates, red asterisks positioned to show the loss or the reduction of SPAG17 isoforms in the mutant tissue (E).

Sanger sequencing of *Pcdo* DNA confirmed the A to T transversion at the start of exon 36, position (c.5236, p.1746; Figure 1B). This mutation introduces a nonsense stop codon (TAA) instead of the wild type lysine (K) amino acid at position 1746. The homozygous *Spag17*^*c.5236A>T*^ mutation segregated completely with the mutant *Pcdo* phenotype and was confirmed by genotyping assays in more than 50 mutant animals over multiple generations to date. Semi-quantitative RT-PCR analysis of *Spag17* RNA using primers spanning exon 33 and 39 indicated high levels of *Spag17* transcript in the testis compared to very limited expression in the lung and brain of 3 months old animals (Figure 1C). Quantification of the band intensity and normalization to a corresponding β-actin band were performed. We found *Spag17* transcript was only marginally reduced in the *Pcdo/Pcdo* mutant tissues as compared to wild-type suggesting the early stop codon did not stimulate a nonsense mediated decay response (Figure 1D). Immunoblotting of the protein lysates extracted from the testis using two SPAG17 antibodies (Figure 1A; Zhang, Jones et al. 2005) showed that translation of all previously reported SPAG17 isoforms (Zhang, Jones et al. 2005) were abolished in the *Spag17*^*Pcdo/Pcdo*^ mutant testis. Our data show a total loss of the SPAG17 75KDa, 97KDa, and the full length 250KDa isoforms in the *Spag17*^*Pcdo/Pcdo*^ mutant testis when compared to wildtype control expression (Figure 1E and data not shown). In the brain, very limited expression of SPAG17 97KDa isoform can be detected but appears at comparable levels between wildtype control and *Spag17*^*Pcdo/Pcdo*^ mutant brains. Similarly, the 250KDa isoform was slightly reduced in the brain tissue lysates of the *Spag17*^*Pcdo/Pcdo*^ animals (Figure 1E). *Spag17* encodes the SPAG17 protein which is known to encode the C1 component of the central pair apparatus of motile cilia (Zhang, Jones et al. 2005). It was previously reported that the *Spag17* germline null animals suffered respiratory distress and died within the first twelve hours of post-natal life due to respiratory motile cilia defects (Teves, Zhang et al. 2013). In the *Spag17*^*Pcdo/Pcdo*^ mutant animals, we observed frequent neonatal death within the first day of postnatal life (17.4%, n=4/23). Mutant animals that survived P0 were viable to weaning age (n>37). *Spag17*^*Pcdo/Pcdo*^ animals that were left to age beyond weaning experienced a shortened life span with pre-mature death around 1 - 4 months of age (n=6/8). Animals were either found dead or were euthanized for humane reasons due to excessive morbidity. This is most likely due to severe hydrocephalus and chronic respiratory insufficiency. Around 25% (n=2/8) of animals lived normally to an age of around 6 months before they were euthanized.

### Normal skeletal development and primary cilia in the *Pcdo* mutants

Previous reports indicated that SPAG17 is required for normal primary cilia function as well as growth and elongation of the long bones in mouse (Teves, Sundaresan et al. 2015). *SPAG17* has also been linked to height in human studies (Weedon and Frayling 2008, Weedon, Lango et al. 2008, N’Diaye, Chen et al. 2011, van der Valk, for the Early Growth Genetics et al. 2014, Wood, Esko et al. 2014). We therefore investigated skeletal development in the *Spag17*^*control*^ (n=16: 3 *Spag17*^*wt/wt*^, 13 *Spag17*^*Pcdo/+*^), and *Spag17*^*Pcdo/Pcdo*^ (n=11) animals. We performed alizarin red and alcian blue skeletal preparations and measured the length of the long bones including the humerus, ulna, radius (Figure 2A&B), femur (Figure 2C&D), and tibia (Figure 2E&F). Statistical analysis (student’s *t-test*) did not detect significant difference in any long bone lengths when comparing data from *Spag17*^*Pcdo/Pcdo*^ mutants to control animals (Figure 2G) except for a slight increase in the *Spag17*^*Pcdo/Pcdo*^ mutant femur length compared to *Sapg17*^*control*^ femur (Figure 2G). The biological significance of this finding remains to be identified. To further assay primary cilia structural development, we generated MEFs from *Spag17*^*Pcdo/+*^ and *Spag17*^*Pcdo/Pcdo*^ animals and performed immunocytochemistry for the ciliary membrane marker, ARL13B. Cilia morphology and occurrence *in vitro* were not disturbed in the *Spag17*^*Pcdo/Pcdo*^ mutant cells compared to control cells (Figure 2H&I). The percentage of ciliated cells was comparable in cells from both genotypes (Figure 2J: n=126 cells from *Spag17*^*control*^ and 155 cells from *Spag17*^*Pcdo/Pcdo*^, 2 animals of each genotype were included in the analysis). These data indicate the *Pcdo* mutation does not affect structure and function of primary cilia in MEFs or long bone development.

**Figure 2:**
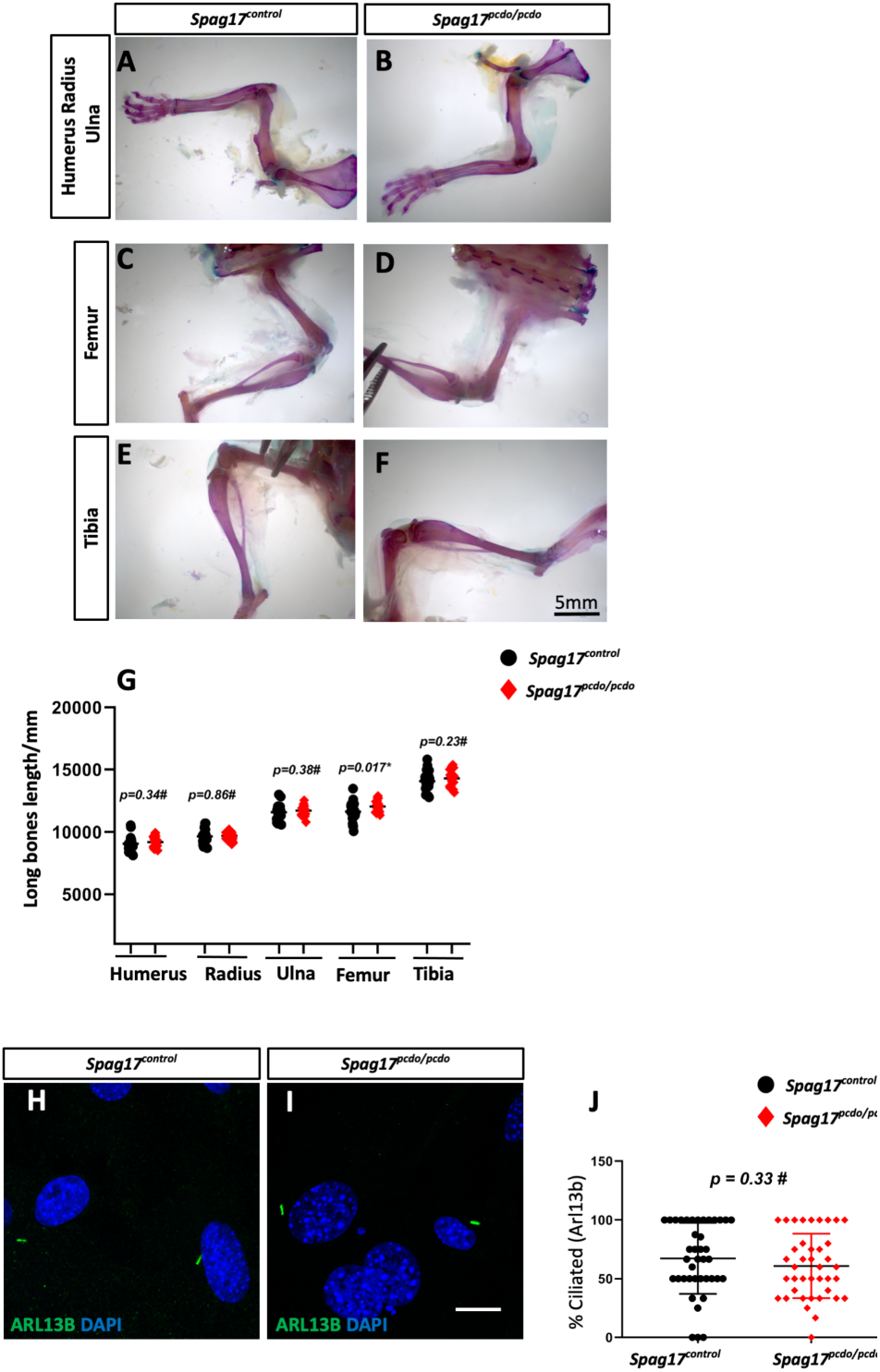
*Spag17*^*Pcdo*^ does not interfere with skeletal development or primary ciliogenesis. Alizarin red and alcian blue stained adult mice upper limb skeleton (A-B), femur (C-D), and tibia (E-F) from animals with the indicated genotypes. Scatter blots of the indicated long bone length, statistical analysis with one *student’s t-test*, did not detect changes in humerus, radius, ulna, femur, and tibia between animals from both genotypes, except for slight increase in the femur length in the *Spag17*^*Pcdo/Pcdo*^ compared to *Spag17*^*control*^ animals, * *p=0,034*, # indicate no significant difference, n= 16, and 11 for *Spag17*^*control*^, and *Spag17*^*Pcdo/Pcdo*^, respectively (G). IHC images of the *Spag17*^*Pcdo/+*^ and *Spag17*^*Pcdo/Pcdo*^ mutant MEFs stained with ARL13B (green), and DAPI (blue) (H - L). Percentage of ciliated MEFs calculated and were not different between *Spag17*^*Pcdo/+*^ and *Spag17*^*Pcdo/Pcdo*^ cells, data obtained from 2 animals each genotype, scatter blot in (J). Scale bars are 5mm (A-F), and10µm (H and I).

### *Spag17*^*Pcdo*^ mice show neonatal progressive hydrocephalus

Hydrocephalus is a prevalent phenotype in PCD mouse models. Similarly, *Spag17*^*Pcdo/Pcdo*^ mutant animals developed a fully penetrant, mildly progressive hydrocephalus. The hydrocephalus was first obvious shortly after birth and before the beginning of the second postnatal week (Figure 3A-F). Measurement of the lateral ventricle surface area from mutants as compared to wild type control animals showed no difference at P0. This measurement was significantly increased in the mutants by P7 and remained significantly higher at P15 and P30 (Figure 3G). No masses of abnormal growth indicative of obstructive hydrocephalus were observed within the ventricular system. However, we did observe severe overt intra cerebro-ventricular and subarachnoid hemorrhage in a small subset of *Spag17*^*Pcdo/Pcdo*^ mutant animals (n=3/32). As expected, this led to blood clots and obstruction of the ventricular system at various points and ultimately severe obstructive hydrocephalus and fatality (Sup. Figure 1).

**Figure 3:**
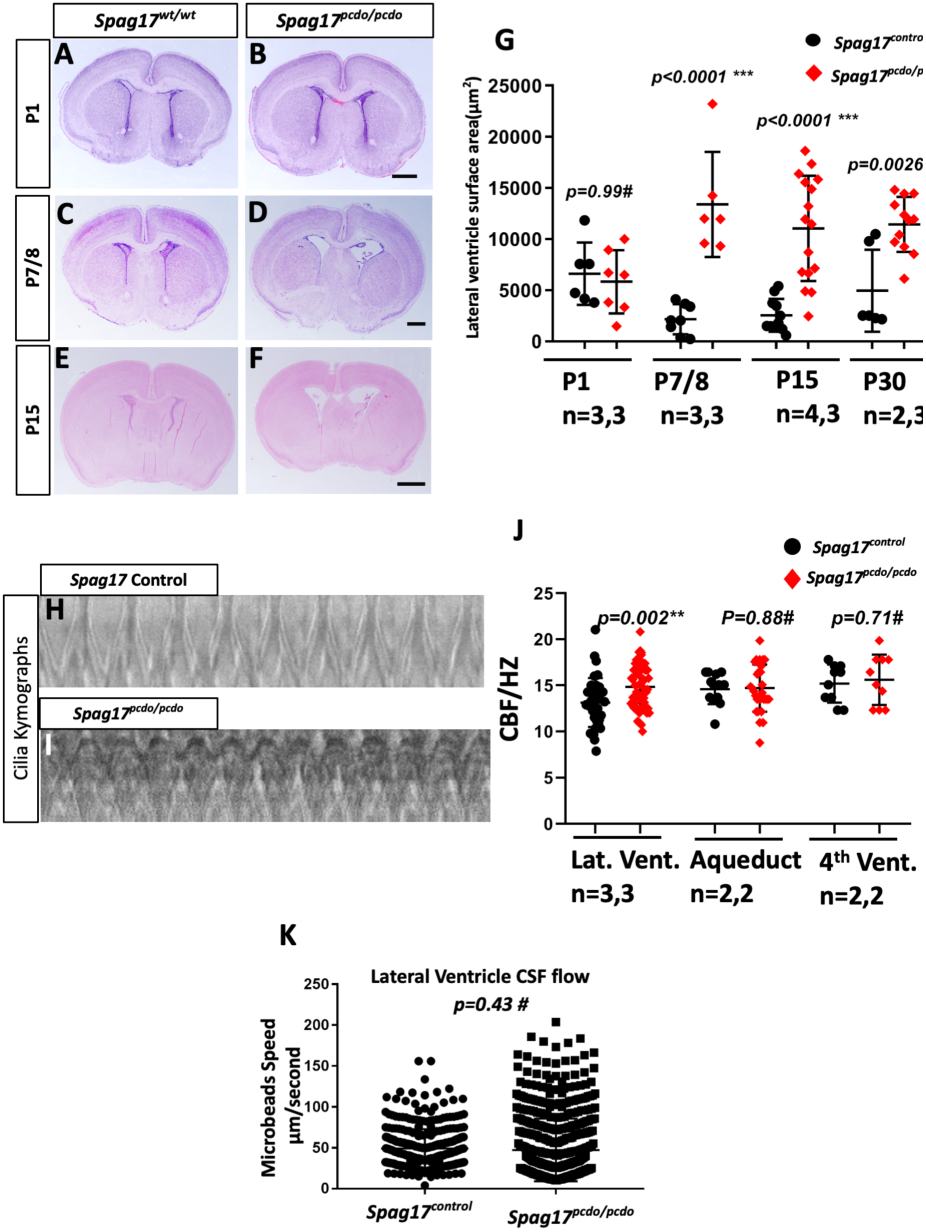
HCP was the first finding distinguished in the *Spag17*^*Pcdo*^ mutant line. Coronal brain sections from *Spag17*^*Pcdo/Pcdo*^ and littermates *Spag17*^*wt/wt*^, showing the dilated lateral ventricles P7 (A-D) mutants pups that continued to be seen in P15 and older animals (E-F). Scatter blots of the lateral ventricle’s surface area measured from animals of the indicated stages of development, no difference at P1 stage, but the lateral ventricle surface area was significantly increased in the *Spag17*^*Pcdo/Pcdo*^ mutants at p7, P15 and P30, data presented were obtained from 3 animals in each genotype and each stage except for *Spag17*^*wt/wt*^ P15 and *Spag17*^*wt/wt*^ P30 n=4 and 2 animals respectively, scale bar (G) Representative kymographs from Spag17^control^ (H) and *Spag17*^*Pcdo/Pcdo*^ (I). Scatter blots of the frequency of the motile cilia beating measurements obtained from lateral ventricle, Aq, and fourth ventricle ependymal cilia. Lateral ventricle cilia are hyperkinetic and beat with a rhythm significantly faster that in the *Spag17* control animals, n=>10 cilia from two animals each genotype (J). No significant difference between speed of the flourescent microbeads flow in lateral ventricle brain slices, n= 4 slice from 2 animals each genotype (K).

The hydrocephalus in the motile cilia models has usually been attributed to perturbations of cilia beating, a highly conserved function of the ependymal motile cilia lining the brain ventricles. This beating function seems essential for directing the cerebrospinal fluid (CSF) towards the next most caudal opening within the cerebro-ventricular system. Given the *Pcdo* phenotypes, we hypothesized ciliary beating may be severely compromised in *Spag17* animals (Teves, Zhang et al. 2013). We examined this using high speed videomicroscoy *ex vivo* at P4 (Sup. video 1& 2). We previously showed that the ependymal motile cilia follow a spatiotemporal pattern of development in the developing forebrain and lateral ventricle (Abdelhamed, Vuong et al. 2018). Ependymal motile ciliogenesis was previously observed along the medial wall of the lateral ventricle around P0 as opposed to the lateral wall, where ependymal ciliogenesis starts towards the end of the first postnatal week (Abdelhamed, Vuong et al. 2018). We also recorded the ciliary beating in other parts of the ventricular system such as the aqueduct and fourth ventricle. Surprisingly, we did not detect any reduction in the ciliary beat frequency in any of the areas we measured (Figure 3H-J). However, we did observe the *Pcdo* mutant motile cilia of the lateral ventricle medial walls were hyperkinetic (Figure 3J). Kymographs of the beating cilia showed restricted waveforms in *Pcdo/Pcdo* mutants (Figure 3H&I), and the mutant cilia were beating at a rate significantly higher than that of the control (p= 0.0016, Figure 3J). Interestingly, kymographs of the beating cilia and analysis of the cilia beat frequency didn’t detect any significant difference in the aqueduct cilia (p=0.88) nor the cilia of the fourth ventricle (p=0.71, Figure 3J). The CSF flow analysis with fluorescent microbeads introduced *ex vivo* into the forebrain slice immediately before the videomicroscopy recording did not show flow speed abnormalities in isolated forebrain lateral ventricle slices (Figure 3K, Sup.video 3& 4). These data are consistent with our conclusion that the *Pcdo* K1746* allele of *Spag17* is a hypomorph as compared to the previously reported *Spag17* null allele which develops paralyzed cilia that affected mucociliary clearance and neonatal survival (Teves, Zhang et al. 2013).

### *Pcdo* aqueductal stenosis disrupts bulk CSF flow

The Aqueduct of Sylvius is a narrow channel for CSF flow connecting the third ventricle to the fourth ventricle. This channel is commonly obstructed in mouse models with deficits in both primary cilia (Town, Breunig et al. 2008) and motile ciliopathy due to collapse and fusion of the walls of the ventricle (Ibañez-Tallon, Pagenstecher et al. 2004). Histological characterization of the aqueduct at P4 from *Pcdo* mutant animals indicated fusion of the ependymal lining at multiple points of the aqueduct (Figure 4A-D). We also noted some enlargement of the sub commissurall organ (SCO) in the *Spag17*^*Pcdo/Pcdo*^ mutant animals (Figure 4E&F). Later in development, the mutant aqueduct appeared collapsed and shrunken (Figure 4G-J). The SCO is an area of highly differentiated ependyma located in the dorso-caudal region of the third ventricle, at the entrance of the Aqueduct of Sylvius, and is well known for secreting high molecular weight glycoproteins necessary for CSF flow and circulation (Rodríguez, Hein et al. 1987). Hydrocephalus is a common feature in animal models with loss or defective development of the SCO (Stoykova, Fritsch et al. 1996, Sakakibara, Nakamura et al. 2002, Blackshear, Graves et al. 2003, Cao and Wu 2015). Collectively, these studies and others suggested that the secretory activity of the SCO is responsible for the maintenance of an open aqueduct. We suspected that aqueductal stenosis could cause localized CSF flow abnormalities and hydrocephalus in the *Spag17*^*Pcdo/Pcdo*^ mutants. We tested this with video microscopy and fluorescent microbeads in the intact aqueductal lumen (Sup. video 5 & 6). We saw that the aqueductal stenosis significantly affected the directionality and speed of the moving beads in the P4 mutant aqueduct (Figure 4K, p<0.00001). We conclude from this data that CSF bulk flow is extremely disturbed due to aqueductal stenosis and that this is what ultimately leads to the hydrocephalus in the *Pcdo* mutants.

**Figure 4:**
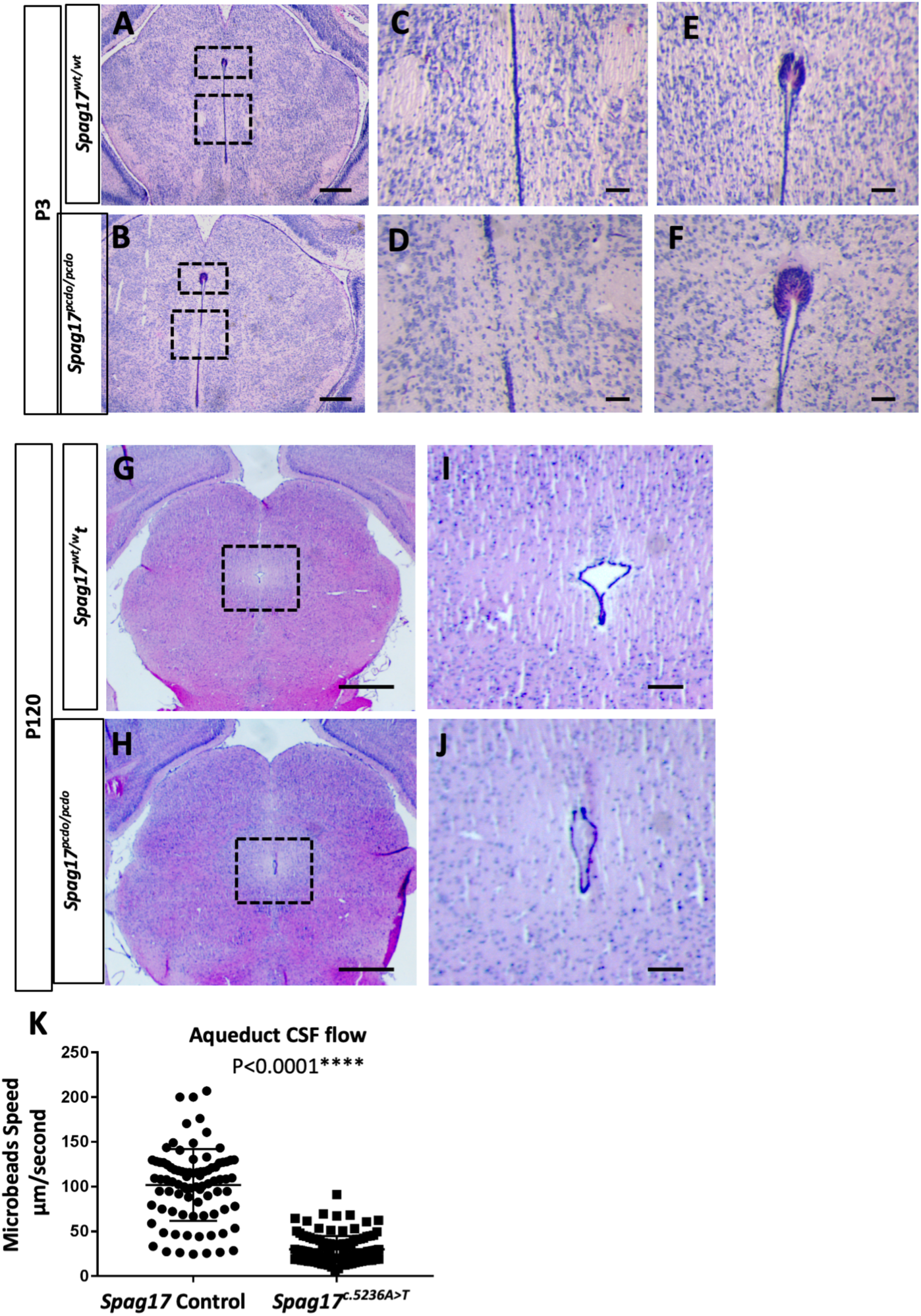
Aqueduct stenosis led to disturbed CSF flow and HCP in the *Spag17*^*Pcdo/Pcdo*^ mice. H&E stained coronal sections through the P3 aqueduct from *Spag17*^*wt/*wt^ (A) and *Spag17*^*Pcdo/Pcdo*^ (B), boxed areas of magnified aqueduct lumen (C and D) and SCO (E and F) and presented to the left. Coronal P120 aqueduct sections from *Spag17*^*+/+*^ (G) and *Spag17*^*Pcdo/Pcdo*^ (H), boxed areas are magnified to the left (I and J). Scale bare is 500µm (A, B, G and H) and 100µm (C, D, E, F, I and J). Scatter blots showing the speed of the moving beads inside the aqueduct, data collected from 4 slices, 2 animals each genotype (K).

### *Pcdo* mutants show other PCD related phenotypes

Gross anatomical assessment of the visceral organs in *Spag17*^*Pcdo/Pcdo*^ mutant showed that organ laterality was unaffected in this model in all animals examined (data not shown). Histological examination of the lung similarly did not show any lung isomerism and gross lung development was not affected in the *Spag17*^*pcd/Pcdo*^ mutants (data not shown). However, detailed histological examination of the lungs and trachea from P8 and P14 *Spag17*^*Pcdo/Pcdo*^ mutants showed abnormal accumulation of homogenous eosinophilic material filling the upper respiratory passages, the trachea and main bronchia indicative of mucus accumulation most likely due to defective mucociliary clearance (Figure 5A-D). Histological investigation of the lungs from older animals showed dilated terminal alveolar ducts (Sup. Figure 2) consistent with chronic obstructive lung disease and bronchiectasis, a predominant finding in PCD patients (Kartagener 1933, Afzelius 1976).

**Figure 5:**
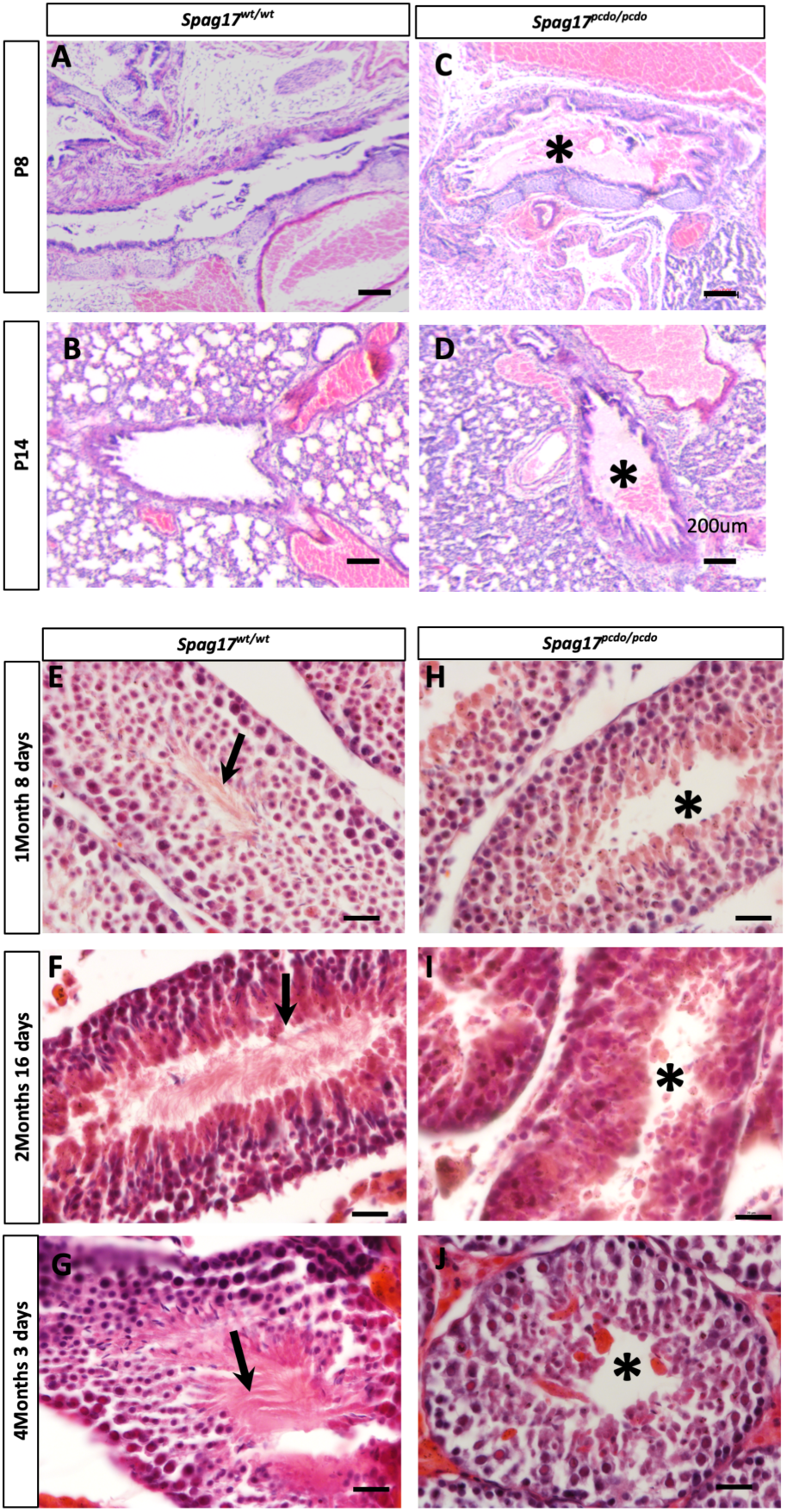
*Spag17*^*Pcdo/Pcdo*^ mutants animals are generally viable and develop phenotypes related to PCD condition. Histology pictures of *Spag17*^*wt/wt*^ and *Spag17*^*Pcdo/Pcdo*^ trachea and lung showing mucus accumulation (asterisks) in the trachea and main bronchus in the mutant animals at the indicted stages (B-E). Seminephrous tubules of the testis from indicated developmental stages and genotypes, *Spag17*^*Pcdo/Pcdo*^ mutant’s lack sperm flagellum at all stages presented, indicted by asterisks in (G, I and K). Arrows in the *Spag17*^*control*^ sections point to mature sperm flagellum (F, H, and J). Scale bar is 200µm (B-E) and 20µm (F-K).

Another common PCD phenotype is male infertility. All male *Spag17*^*Pcdo/Pcdo*^ animals (n= 4) we tested were infertile and were unable to produce any litters when mated with wild type control female mice (when paired twice for two weeks each attempt, n=8). Conventional histological analysis of the testis was performed at 5 weeks, 10 weeks, and 4 months of age. In all stages of testicular development we examined, the *Spag17*^*pcd/Pcdo*^ mutant seminiferous tubules were devoid of any sperm with a clear lumen while all control sections showed evidence of normal spermatogenesis with mature sperm observed in the center of the seminiferous tubules (Figure 5E-J). No clear difference is noted at the level of the germ cells or the developing spermatocytes number nor morphology (Figure 5E-J). The round spermatids failed to fully mature and failed to produce flagella (Figure 5E-J). This indicates that spermiogenesis process including sperm flagellum development requires SPAG17.

### Differential requirements of *Spag17* for motile ciliogenesis in the brain, lung, and sperm flagellum

Given the functional deficits we observed in the motile cilia, we performed a morphological analysis of the motile ciliary structure. We first performed immunohistochemical analysis with the acetylated α-tubulin antibody to stain the ciliary axoneme. No apparent morphological differences were detected in the *Spag17*^*Pcdo*^ mutant ependymal and/or respiratory cilia when compared to wildtype control cilia (Figure 6A-D). We saw similar results with a more stringent scanning electron microscopy analysis. The abundance of cilia and cilia length appeared comparable in ependymal cilia lining the medial and lateral walls of the forebrain lateral ventricle indicating that the *Spag17*^*Pcdo*^ allele does not perturb production of motile cilia in the brain ependymal cells or the respiratory epithelial cells (Figure 6E-H). In contrast, *Spag17*^*Pcdo/Pcdo*^ mice showed deficits in sperm flagellum formation. Consistent with the histological analysis above, immunohistochemical staining of testicular sections from *Spag17*^*Pcdo/Pcdo*^ mutants showed a total lack of acetylated α-tubulin staining at 5 and 10 weeks of age as compared to the age matched control where we observed clear and pronounced acetylated α-tubulin staining (Figure 6I-L).

**Figure 6:**
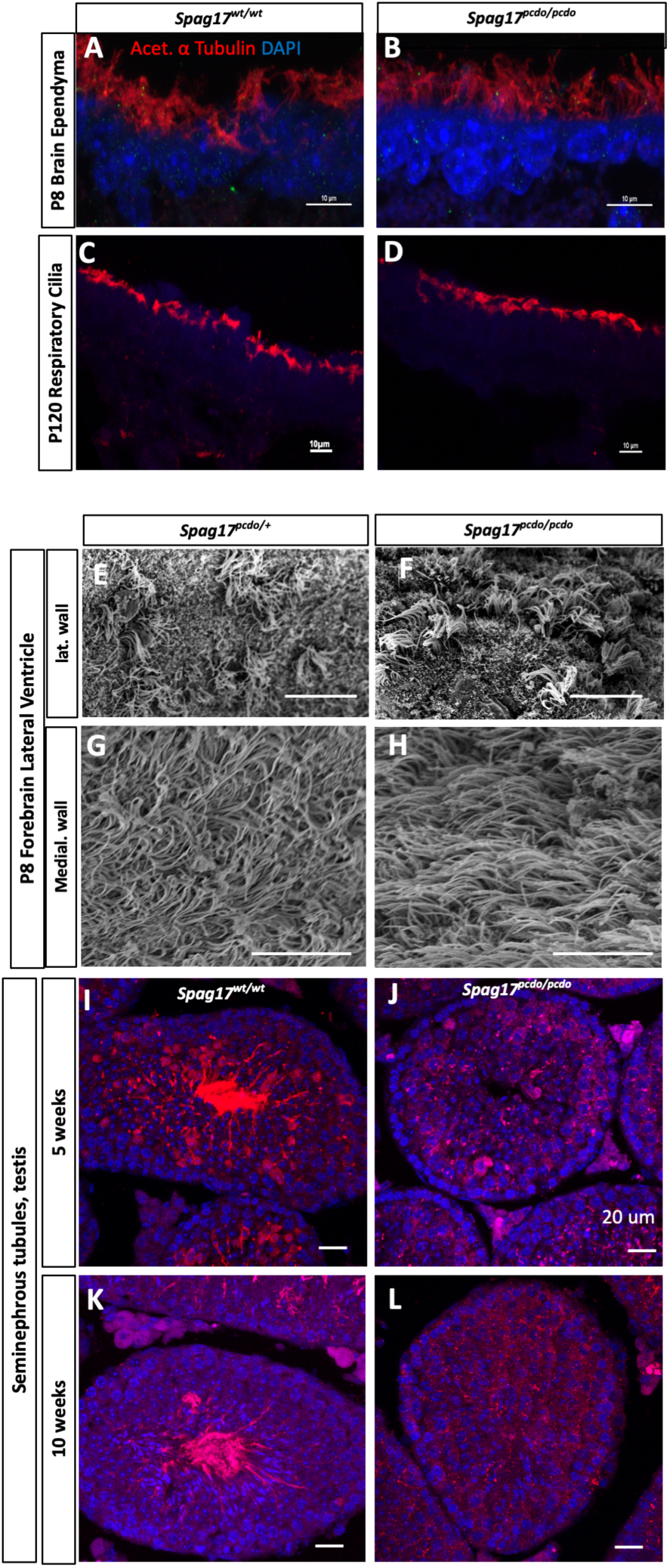
SPAG17 is essential for sperm flagellum development but not for initiation of motile ciliogenesis in the brain or the respiratory tract. IHC images of forebrain brain ependymal (A and B), tracheal epithelial (C and D) cells of the indicated genotypes and stage stained for cilia axonme with Acet.α.Tubulin (red) and DAPI to stain nuclei (blue). SEM images of the lateral wall (E and F), medial wall (G and H) of the P8 forebrain ependymal cilia. No clear difference in the overall morphology of the ependymal or the respiratory motile cilia. IHC images of the seminephrous tubules stained for Acet.α. Tubulin (red) and DAPI (blue). No cilia staining can be detected in the *Spag17*^*Pcdo/Pcdo*^ mutant seminephrous tubules compared to *Spag17*^*wt/*wt^ (I-L). Scale bars are 10µm (A -H), 20µm (I-L).

### Distinct ultrastructural defects in the *Spag17*^*Pcdo*^ mutant respiratory and brain ependymal motile cilia

SPAG17 is a cilia structural protein essential for development of the C1a process of C1 microtubule structure of the central pair apparatus (Zhang, Jones et al. 2005). We therefore investigated the ultrastructure of the ependymal and respiratory motile cilia using transmission electron microscopy. Ependymal cilia cross sections appeared fairly normal with no clear defects in either the structure or the orientation of the central pair apparatus or other motility structure such as inner or outer dynein arms (Figure 7A-C). This is consistent with the limited functional disturbance in ependymal cilia beat frequency and the undisturbed localized (intraventricular) CSF flow observed in this model. In contrast to above mentioned defects in the ependymal cilia, TEM analysis of the respiratory cilia cross sections showed predominantly abnormal respiratory tracheal cilia in the *Spag17*^*Pcdo*^ mutants. Two types of related deformities can be clearly observed. Firstly, mutant respiratory cilia lack one of the central pair microtubule singlets in a subset of the mutant cilia with appearance of 9+1 microtubule organization (Figure 7D-G). Most of the remaining cilia in the mutant trachea have a distorted central pair apparatus compared to wildtype control animals (Figure 7H-K). Secondly, 22.2% of the *Spag17*^*Pcdo*^ mutant central pairs (n=43) of respiratory cilia were not equally aligned, and seen acquiring disorganized angles or in opposed orientation compared to only 16.3% of wild type cilia (n=18) (Figure 7L&M). This indicates that rotational planar cell polarity of the central pair was perturbed in the *Spag17*^*Pcdo/Pcdo*^ mutant respiratory cilia. Orientation of the axonemal central pair of microtubules was measured by drawing a line through the central pair and measuring the angle of the line with respect to the 0–180° angle arbitrarily set for the first line drawn, as previously described (Kunimoto, Yamazaki et al. 2012, Hegan, Ostertag et al. 2015). The deviation from this angle is significantly greater in the *Pcdo* mutant cilia. This indicates that the SPAG17 is not only essential for central pair development but could also be indispensable for establishing the central pair rotational polarity. The observed degree of central pair disorientation could have a strong influence on the highly coordinated ciliary beating function and thereby contribute to impaired directional fluid flow and a mucociliary clearance defect in the lung.

**Figure 7:**
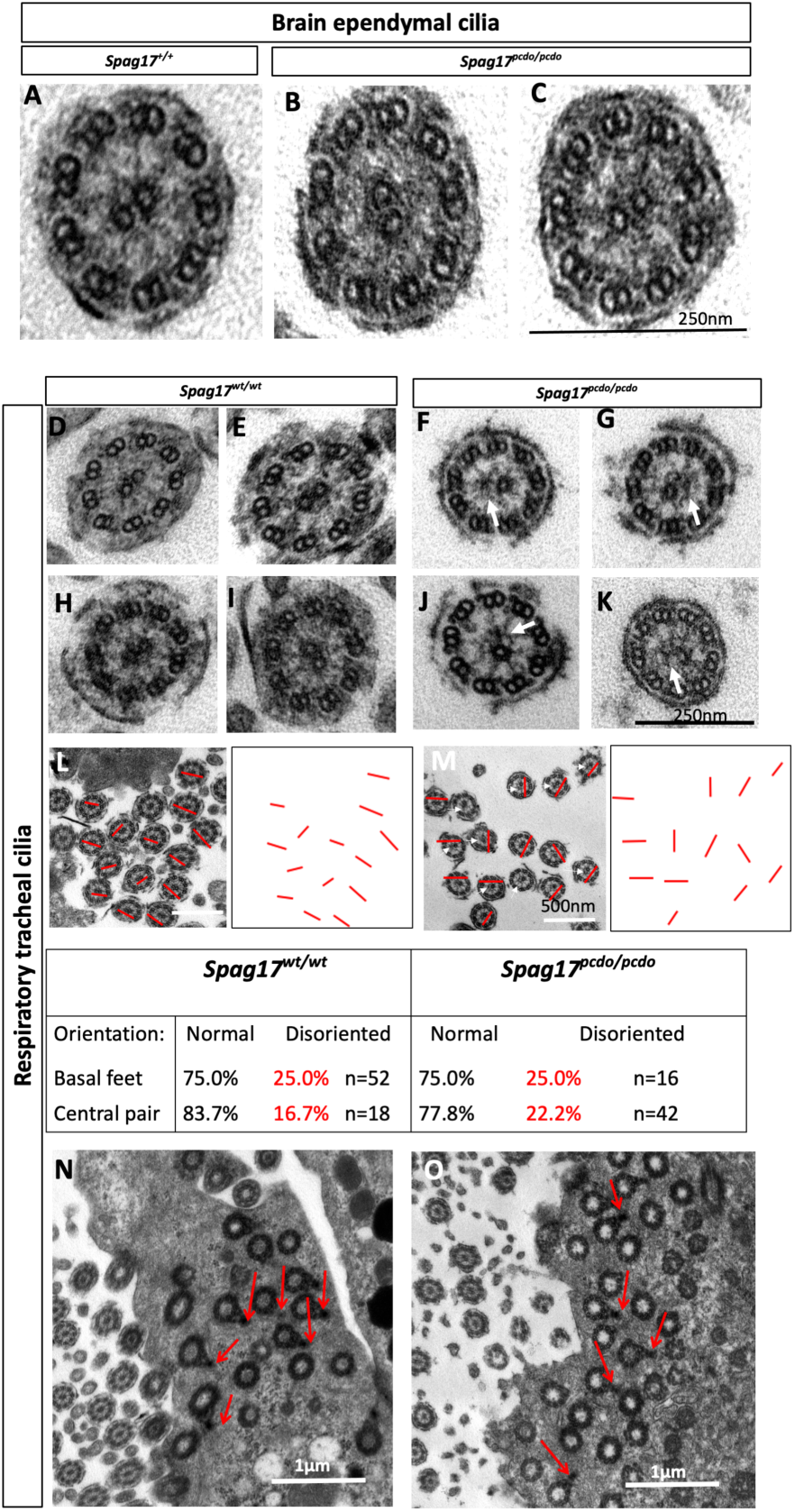
SPAG17 is essential for central pair C1 microtubule development and stability in the respiratory epithelial cilia. TEM cross section of the ependymal cilia from *Spag17*^*wt/wt*^ and *Spag17*^*Pcdo/Pcdo*^ showing no clear ultrastructural defects in the mutant ependymal cilia (A-C). Higher magnification of the respiratory cilia axonem cross sections showing lack of (Arrows in F and G), or distorted abnormal (arrows in J and K) C1 microtubule structure of the central pair apparatus in the *Spag17*^*Pcdo/Pcdo*^ mutants compared to wildtype respiratory cilia cross sections (C, D, G and H). TEM images of the respiratory epithelium of *Spag17*^*wt/wt*^ and *Spag17*^*Pcdo/Pcdo*^ (L and M) animals, red lines indicate the central pair axis of orientation in sections from both genotypes, the *Spag17*^*Pcdo/Pcdo*^ mutant central pair(s) were abnormally oriented in different directions, blue thin arrows in (M) point to cilia cross section that have no C1 microtubule. Basal processes of the basal bodies were normally and equally aligned in the respiratory epithelial cells from *Spag17*^*wt/wt*^ and *Spag17*^*Pcdo/Pcdo*^ mutants (N and O, red arrows). Scale bars are 250nm (A and K), 500nm (L and M), and 1µm (N and O).

The observed central pair orientation defect prompted us to also assay the orientation of the basal body basal feet in the mutant respiratory cilia as compared to wildtype. The directionality of ciliary beating is determined by the orientation of basal feet of the basal bodies, which are usually positioned at the fourth, fifth and sixth circularly arranged triplets of microtubules within the basal bodies. The basal feet associated with the basal bodies are usually oriented in the direction of the effective stroke of the ciliary beating and this is controlled by planar cell polarity pathway (Gibbons 1961, Boisvieux-Ulrich, Laine et al. 1985, Kunimoto, Yamazaki et al. 2012). However, we found that in the *Spag17*^*Pcdo*^ mutant cilia (n=16 cilia), the basal feet of the basal body appeared normally aligned when compared to wildtype control cilia (n=52 cilia, Figure 7N&O). We did not observe further defects in other structures required for ciliary motility such as inner and outer dynein arms.

Altogether, our data indicate there is differential requirement for SPAG17 in different cell types which produce motile cilia. We conclude SPAG17 is critical for sperm flagellum initiation and formation while it has a more specific role (limited to central pair apparatus and C1a projection development and orientation) during respiratory cilia structural development and an even more limited role during cilia differentiation and maturation of the brain ependymal cells. We also conclude the *Spag17*^*Pcdo*^ allele has no effect on the structural or functional development of the primary cilia.

## Discussion

### *Pcdo* is an allele of *Spag17* revealing tissue specific effects on SPAG17 translation

In this study, we present a detailed characterization of the *Pcdo* mutant we recovered from an ENU forward genetic screen and determined to be a nonsense allele of *Spag17* (Figure 1A&B). The *Pcdo* variant is a c.5236A>T nonsense mutation predicted to introduce premature stop codon at position 1746 of the 2320 amino acid SPAG17 protein. Significantly higher levels of *Spag17* RNA were expressed in the testis as compared to the lungs and brain tissues of the adult mice. This data is consistent with previous *Spag17* expression analyses (Zhang, Jones et al. 2005). The *Pcdo* mutation resulted in reduced RNA expression that was not statistically significant between the wild-type and mutant tissues (Figure 1C&D), indicating that *Pcdo* mutation does not induce nonsense mediated decay in any tissue examined.

Protein analyses of mutant tissue samples from testis and brain indicated that *Pcdo* mutation leads to the loss of SPAG17 protein isoforms in testis but not in brain, although the ENU-induced variant is common to full length isoform of *Spag17* known to be expressed in testis, brain, and lung. However, our analysis also identified very low levels of the SPAG17 97KDa isoform in the brain which did not seem to be affected in the *Pcdo* mutants. Similar observations have been documented in studies of another motile cilia gene, *adenylate kinase 7*(*Ak7*), in which mutation leads to loss of protein in the testis but not in the respiratory cilia (Lorès, Coutton et al. 2018). Our data indicate that proteins expressed in both motile cilia and sperm flagella may have tissue specific properties in each of these two highly related organelles. This may be attributed to the major differences between sperm flagella and simple motile cilia in the brain and the respiratory epithelium.

### Mild phenotypes observed in *Pcdo* mutant animals indicate that *Pcdo* is a hypomorphic allele of *Spag17*

*Spag17*^*Pcdo/Pcdo*^ mice developed early postnatal communicating hydrocephalus, have mucus accumulation in the respiratory passages and were infertile. Although the majority of *Pcdo* mutant animals survived weaning, they have a shortened life span as compared to controls. We attribute this to chronic morbidity. All these phenotypes are consistent with motile cilia involvement in the *Pcdo* mutants. In spite of these indicators of motile cilia deficits, beating and fluid flow were not altered in the mutant lateral ventricle. Rather, we determined the cause of the hydrocephalus to be progressive aqueductal stenosis affecting the CSF fluid turbulence and ventricular bulk flow. *Pcdo* mutant mice did not have obvious primary cilia form or function defects. These findings that *Pcdo* mutants develop milder gross anatomical and ciliary motility phenotypes and survive longer than the *Spag17* knockout mouse phenotypes (Teves, Zhang et al. 2013) support our conclusion that *Pcdo* is a hypomorphic allele of *Spag17*.

### Motile and primary ciliopathies can overlap in the same model

Motile and primary ciliopathies have long been considered as two distinct dysfunctions of two related but functionally different organelles, the motile and non-motile cilia, respectively. However, recent advances and more characterization of ciliopathy mutants indicated that motile and primary ciliopathies can overlap in the same animal model and/or human patients (Moore, Escudier et al. 2006, Fliegauf, Benzing et al. 2007, Ferkol and Leigh 2012, Bukowy-Bieryłło, Ziętkiewicz et al. 2013). Characterization of the *Spag17* knock out mice provided new evidence that motile and primary ciliopathies can overlap or develop in the same model organism (Teves, Zhang et al. 2013),(Teves, Sundaresan et al. 2015). *SPAG17* mutations are reported to cause human PCD (Andjelkovic, Minic et al. 2018) and variants in the gene have been linked to human height (Weedon and Frayling 2008, Weedon, Lango et al. 2008, Takeuchi, Nabika et al. 2009, Kim, Lee et al. 2010, Zhao, Li et al. 2010) and multiple goat body measurements (Zhang, Jiang et al. 2019). Interestingly, combined missense changes in *SPAG17* and *WDR35* causes complex neurodevelopmental malformations and skeletal cranioectodermal dysplasia, a primary ciliopathy (Córdova-Fletes, Becerra-Solano et al. 2018). These studies altogether confirm that *Spag17* has a role in development and function of both motile and immotile primary cilia, but the effects differ based on the precise allele in question.

### Similar to other mutants in central pair genes, *Pcdo* mutants exhibited no beating abnormalities and have no laterality defects

A hallmark of a central pair PCD ciliopathy is the absence of laterality defects, subtle ciliary beating abnormalities, and unequivocal ultrastructural defects of the ciliary axoneme. Vertebrate organ asymmetry is well known to require the nodal cilia that generate leftward flow across the node that leads to asymmetric distribution of morphogens in the developing embryo. These cilia are motile because they express inner and outer dynein arms but do not have a central pair apparatus. It is therefore intriguing to see almost all central pair protein mutants, including this allele of *Spag17*, do not develop laterality defects (Davy and Robinson 2003, Lechtreck, Delmotte et al. 2008, Cindrić, Dougherty et al.). However, mutants of the dynein complexes consistently develop various degrees of laterality defects including dextrocardia and heterotaxy (Bartoloni, Blouin et al. 2002, Dougherty, Loges et al. 2016, Abdelhamed, Vuong et al. 2018, Loges, Antony et al. 2018, Ta-Shma, Hjeij et al. 2018). Lack of laterality in the *Spag17*^*Pcdo/Pcdo*^ mutants further support the model that central pair proteins including *Spag17* are dispensable for organ asymmetry and body laterality development.

### Ependymal cilia show normal ultrastructure and minimal beating defects

The main function of the ependymal cilia is to beat and generate cilia-directed CSF flow. The forebrain ependymal cilia in the *Spag17*^*Pcdo*^ mutant have no apparent ultrastructural abnormalities and the cilia beating frequency was unaffected in the ependymal cilia of the aqueduct and cilia of the fourth ventricle ependymal (Figure 3K). Surprisingly, a subset of the ependymal cilia on the medial wall of the lateral ventricle in the *Spag17*^*Pcdo/Pcdo*^ mutants are hyperkinetic and beat with a rate significantly higher than that of the control cilia (Figure 3J). The hyperkinetic beating does not disrupt the local cilia mediated flow in the acute *ex vivo* isolated medial wall forebrain slice (Figure 3K). It is possible that this could have a prominent effect *in vivo* due to more limiting space constraints. Alternatively, the lack of an effect on the ependymal cilia structural development and function could be due to the sufficient expression of near normal levels of SPAG17 proteins in brain ependymal cells, which we demonstrated with immunoblotting. Interestingly, the hyperkinetic cilia phenotype in the *Spag17*^*Pcdo/Pcdo*^ mutants is in accordance with other motile cilia mutants that have normal motile cilia ultrastructure but hyperkinetic cilia such as *DNAH11* PCD patients (Bartoloni, Blouin et al. 2002, Schwabe, Hoffmann et al. 2008, Pifferi, Michelucci et al. 2010, Knowles, Leigh et al. 2012, Dougherty, Loges et al. 2016) and a mouse model(Handel and Kennedy 1984). Assessment of this beating pattern on the CSF flow *in vivo* would be a better indicator of the effect of the hyperkinetic cilia on the fluid flow and hydrocephalus development. This will be helpful to further explain if the hyperkinetic cilia phenotype is directly causing the hydrocephalus in this model or other mechanisms are responsible.

### The hydrocephalus in the *Spag17*^*Pcdo/Pcdo*^ mutants is due to aqueductal stenosis

The Aqueduct of Sylvius in the *Pcdo* mutants appeared stenotic with collapsed walls and was associated with a slight enlargement of the SCO (Figure 4). In other models, patency of the aqueduct is compromised in the presence of dysfunctional SCO or SCO ependymal cilia (Pérez-Fígares, Jimenez et al. 2001, Swiderski, Agassandian et al. 2012), leading to aqueductal stenosis or occlusion and non-communicating hydrocephalus. Ciliary beating function and directionality of the beating were defective when assayed in intact aqueduct lumen. The stenosis could have led to a mechanical constraint on the ciliary beating and thus to significantly reduced flow of CSF into the fourth ventricle. We concluded that *Pcdo* mutants have communicating hydrocephalus as the aqueduct was stenotic and not completely occluded indicative of a potential milder dysfunction of the SCO.

Surprisingly, we noted severely defective respiratory motile cilia structure consistent with a central pair protein insult. The *Spag17*^*Pcdo*^ mutant cilia either lack one of the central microtubule or have distorted central pair structure. The orientation of the central pair also appeared to not align in one direction indicative of disorganization of cilia central pair polarity (Figure 7 L and M). This is expected to interfere with a directional fluid flow in the respiratory passages and lead to mucus accumulation and recurrent infection. The axis of polarity is controlled by signaling and mechanical cues. The mechanical cue is a cilia-generated fluid flow (Mitchell, Jacobs et al. 2007). The intense mucus accumulation observed in the *Spag17*^*Pcdo*^ mutants indicate that ciliary motility in the lung is defective. This is consistent with the structural defects and seems to be the major cause of morbidity and the shortened life span in these mutants. Moreover, sperm flagellum development was completely inhibited in the *Pcdo* mutants and no mature spermatozoa with flagellum can be detected at any postnatal stage of testicular development (Figure 5E-J & Figure 6I-L).

Altogether, our data indicate that SPAG17 is necessary for sperm flagellum development, important for proper assembly of central pair apparatus in the respiratory motile cilia but does not have a similar requirement in ependymal cilia development. *Spag17* is highly expressed in testis, has moderate expression levels in the lungs, and lowest levels were observed in the brain (Figure 1C&D). It seems like there is a correlation between the basal *Spag17* expression levels and the downstream results of *Pcdo* mutation in the different motile cilia examined. Spermatocytes and round spermatids seem to absolutely require SPAG17. While other motile cilia cell types can be more tolerant to *Pcdo* mutation, the respiratory cilia cannot compensate for the mild reduction in the *Spag17*. At the protein level, SPAG17 is known to interact with other central pair proteins such as SPAG16 and SPAG6 and to form a complex at the C1a microtubule projections (Wargo, Dymek et al. 2005, Zhang, Jones et al. 2005, Fu, Zhao et al. 2019, Zhao, Hou et al. 2019). One explanation of this finding is that the minimal levels of the SPAG17 produced in the *Pcdo* mutant ependymal cells was sufficient to facilitate the interaction with other proteins and to form C1a projection but much more of SPAG17 was needed in the respiratory cilia to form these protein complexes and thereby form the correct central pairs of the motile respiratory cilia. All isoforms of SPAG17 were completely abolished in the *Pcdo* mutant testis and so it seems that all C1a projection interactome were unable to form and this led to complete failure of sperm flagellum formation. Several testis specific proteasome/ubiquitin enzymes have been reported in rodents (Mochida, Tres et al. 2000) (Nishito, Hasegawa et al. 2006) and human(Mitchell, Woods et al. 1991, Uechi, Hamazaki et al. 2014). It is also possible that the mutant SPAG17 protein is degraded in the testis by a sperm specific proteasome-mediated quality control mechanism (Morales, Kong et al. 2003, Zimmerman and Sutovsky 2009, Sutovsky 2011). Altogether, these data are consistent with recent reports that effects of specific gene mutations on the protein translation and thereby function may dramatically differ between various cell types expressing the same allele (Lucas, Davis et al. 2019).

In summary, this study describes a novel allele of *Spag17* that encodes the C1a projection SPAG17 protein. The *Pcdo* allele recapitulated most PCD phenotypes previously observed in other models due to central pair abnormalities. These are often the most difficult to diagnose in human due to lack of laterality phenotypes, only subtle beating defects and largely undisturbed cilia ultrastructure (Edelbusch, Cindrić et al. 2017). We suggest *Spag17*^*Pcdo*^ is very useful model to study the pathogenesis and molecular mechanisms employed in the development of human PCD subsequent to central pair defects. Further analysis of the *Pcdo* mutant line could help uncover a new diagnostic method of central pair PCD human condition.

## Material and methods

### Ethyl-N-nitrosourea (ENU) mutagenesis and recovery of *Pcdo* mutants

ENU mutagenesis was performed as described (Herron, Lu et al. 2002, Stottmann, Moran et al. 2011). Briefly, 6 to 8-week old C57BL/6 males were injected intra-peritoneally route with a weekly fractioned dose of ENU to produce G0 males which were bred to FVB females (Jackson laboratories) to generate G1 heterozygous carrier males. These G1 males were then outcrossed to FVB females to generate G2 potentially heterozygous carrier females. These G2 females were backcrossed to their respective G1 male parent to generate G3 embryos or pups which were screened for organogenesis phenotypes. a mutation of interest that could reproducibly produce developmental defects. *Pcdo* mouse were first identified with the dilated brain ventricles and HCP. All protocols were approved by the Cincinnati Children’s Hospital Medical Center IACUC committee (IACUC2016-0098).

### Histological analysis

Neonatal pups were sacrificed through decapitation and for adult histology, littermate animals underwent cardiac perfusion using cold heparinized phosphate buffered saline (PBS) and formalin (SIGMA) solution. Brains, lungs and testis were dissected and fixed for up to 72 h in formalin at room temperature followed by immersion in 70% ethanol (for histology). Samples were then paraffin embedded and sectioned at 6µm thickness. Sections were processed for hematoxylin and eosin (H&E) staining with standard methods. Histology sections were imaged using a Zeiss Discovery.V8 Stereoscope or Nikon NIE upright microscope.

### Immunohistochemistry (IHC)

IHC were performed as previously described (Driver, Shumrick et al. 2017). Briefly, fixed paraffin blocks from control and mutant animals were sectioned at a thickness of 6µm. Sections were de-paraffinized, rehydrated in graded ethanol, antigen retrieval were performed by boiling in citrate buffer pH 6,00 for 1 minute in a microwave. Nonspecific antigens in the tissue sections were blocked in 4% normal goat serum in PBS-tween and the anti-acetylated α tubulin (1/2000, Sigma # T6793) antibody was incubated with the tissue overnight at 4°C, followed by 3 stringent washes in PBS-tween and application of Alexa flour® 596 conjugated goat anti mouse secondary antibody (A11005, Invitrogen) for 1 hour followed by 3 washes in PBS tween. Nuclei were counter stained with 4′,6-diamidino-2-phenylindole (DAPI) and mounted in ProLongTM Gold antifade mounting media (Invitrogen). Images were captured using Nikon C2 confocal microscope.

### Whole exome sequencing

Whole exome sequencing was done with the BGI Americas Corporation using standard protocols.

### Sanger Sequencing

The *Spag17* nonsense variant in exon thirty six identified by whole exome sequencing was further confirmed by Sanger sequencing using the following forward primer: TTTGCCATTTGGATTTACAGG and reverse primer: GTAGCACTTTGGGTCCTTCG. PCR amplicons from *Spag17*^*+/+*^, *Spag17*^*Pcdo/+*,^ and *Spag17*^*Pcdo/Pcdo*^ animals were purified and concentrated using the Zymo DNA clean and Concentrator kit (Zymo Research Corporation, Irvine, CA). 100ng of concentrated DNA were used in each Sanger sequencing reaction.

### *Spag17*^*Pcdo*^ mouse line genotyping assays

All animals and pups obtained from the *Spag17*^*Pcdo*^ mouse line were genotyped using the TaqMan® Sample-to-SNP™ kit (Applied Biosystems) for a single-nucleotide change at mouse chr1:g,100088282A>T (assay ID ANWCWCA). *Spag17*^*Pcdo*^TaqMan® Sample-to-SNP™ assays were performed according to manufacture instruction and run using QuantStudio 6 Real Time PCR machine (Applied Biosystems).

### Real time semi-quantitive PCR (RT-PCR)

Total RNA was extracted from testis, lungs, and whole brains of P30 mice using Trizol^®^ reagent (Thermofishers cientific) according to manufacturer instructions. 5µg of total RNA was used for reverse transcription into cDNA using the SuperScript III First strand synthesis system (Invitrogen) according to manufacturer directions. Final cDNA products were diluted 1:10 and then 2µl of the diluted cDNA were used to amplify *Spag17* transcript using primers that span exon 33 and exon 39 (Forward primer: GATGGAGGGCTACGAAAGC and reverse primer: AACTGTTAGGTGGGCTGCAA.) β-actin was used as loading control (forward primer: GTGACGTTGACATCCGTAAAGA,reverse primer: GCCGGACTCATCGTACTCC).

### Western immunoblotting

P33 mouse brain, and testis tissues were lysed in Pierce RIPA buffer (Thermo #89901) containing protease Inhibitor cocktail (Roche #11697498001). Protein concentration in the whole cell extracts was determined with the BCA colorimetric assay (Thermofisher) according to manufacturer instructions. Denatured proteins were separated by electrophoresis on a gradient 4-12% Tris-glycine gel. Protein was transferred to a PVDF membrane, blocked in Odyssey blocking buffer and incubated over night at 4°C with 1:3000 rabbit anti-N. terminus or c.terminus anti-SPAG17 antibodies (previously described by Zhang and colleague, Zhang, Jones et al. 2005) and 1:1000 Mouse anti-Tubulin (Sigma #T6199) antibodies). Membranes were washed and incubated for 1 hour in 1:15000 goat anti-rabbit IRDye 680CW (LICOR) and 1:15000 goat anti-mouse IRDye 800Rd (LICOR) and bands were visualized on LICOR Odyssey imaging system.

### Scanning electron microscopy (SEM)

The medial walls and lateral walls of the P4 forebrains were processed for SEM as previously described (Abdelhamed, Vuong et al. 2018). Briefly, samples were fixed in electron microscopy grade 2% PFA and 2.5% glutaraldehyde in 0.1 M Na cacodylate buffer (pH 7.4) at 4°C overnight. Tissues were washed thoroughly in 0.1 M Na cacodylate buffer (pH 7.4) then post-fixed in 1% osmium oxide (diluted in 0.1 M Na cacodylate buffer) for 1 hour. Samples were washed thoroughly in 0.1 M Na cacodylate buffer and dehydrated prior to critical point drying in 100% ethanol. Brain tissues were then coated with gold palladium using a sputter coater (Leica EM ACE600) and scanned with a Hitachi SU8O1O scanning electron microscope.

### Transmission electron microscopy (TEM)

Brain ependymal cells and tracheal epithelial cells were fixated in 2.5% glutaraldehyde and processed for TEM analyses by standardized protocols as previously reported (Wallmeier, Frank et al. 2019). Sections were collected on copper grids, stained with Reynold’s lead citrate and visualized using the Philips CM10 or Jeol 1400+.

### High -Speed videomicroscoy of cilia, cilia beat frequency, and CSF flow analysis

High speed videomicroscoy of the beating ependymal cilia, cilia beat frequency and CSF flow analysis were performed as previously described (Abdelhamed, Vuong et al. 2018). Briefly, brains were dissected out from P4 *Spag17*^*Pcdo/Pcdo*^ or wiltype control animals in Dulbecco’s Modified Eagle Medium: Nutrient Mixture F-12 (DMEM/F12) supplemented with L-glutamine (Gibco) and 1% N2 supplement (Gibco) at room temperature, and serial 200 µm-thick coronalsection. Sections were obtained from the forebrain and aqueduct. Areas from medial wall of the lateral ventricle were further microdissected and subjected to video microscopy recording. For concurrent imaging of the beating cilia with green fluorescent micro-beads (FluoSpheres, Thermo Fisher) were introduced into slices immediately before video recording and we used an inverted Nikon Ti-E wide-field microscope fitted with a Nikon 40× Plan Apo 0.95 N.A. air objective and an Andor Zyla 4.2 PLUS sCMOS monochromatic camera. Light was channeled through a custom quad-pass filter and 300 frames were collected at ∼40 frames/s. Tracking and quantitative analysis of the moving beads were performed using NIS Elements software. Beads tracking and cilia beat frequency analysis were performed using NIS Element software, version 4.1. Statistical analysis and pairwise comparison performed using GraphPad Prism software.

### Skeletal preparations

For skeletal preparations, P30 animals were sacrificed by high dose of anesthesia using isoflurane followed by trans-diaphragmatic cardiac extrusion. Animals were eviscerated and fixed for 2 days in 95% ethanol. They were stained overnight at room temperature in Alcian blue solution (Sigma #A3157) containing 20% glacial acetic acid. Excess stain was cleared in 95% ethanol for 24 hours and skeletons were then slightly cleared in a 1% KOH solution overnight at room temperature. They were then stained overnight in Alazarin red solution (Sigma #A5533) containing 1% KOH. A second round of clearing performed by incubating tissues in 20% glycerol/1%KOH solution for 24 hours. Finally, they were transferred to 50% glycerol/50% ethanol for photography. Skeletal preparations were imaged using a Zeiss Discovery.V8 Stereoscope.

### Generation of MEFs Culture

MEFs were generated from E13.5 embryos. Embryos were dissected in PBS, decapitated, and eviscerated. The remaining tissue was incubated in trypsin overnight at 4°C. Tissue fragments were incubated with the trypsin in 5%CO2 incubator for 30 minutes. Cells were then allowed to grow to confluency in complete DMEM containing 10%FBS and penicillin/streptomycin. MEFs were stained within three passages of their isolation. Cells were stained for ARL13B (ProteinTech # 17711-1-AP) (1/500), ciliary membrane marker and DAPI to stain nuclei.

## Acknowledgements

We thank Chelsea Menke for technical assistance throughout this project, Maria Teves for helpful discussion and the SPAG17 antibody.

