## Supplemental materials for "A novel hypomorphic allele of *Spag17* causes primary ciliary dyskinesia phenotypes in mice"

**Supplemental Material**

**Supplemental Figure 1:** Overt severe subarachnoid and intraventricular hemorrhage occurs in less than 10% of the weaned *Spag17*$^{Pcdo/Pcdo}$ mutants, but is never seen in *Spag17*$^{+/+}$ animals.

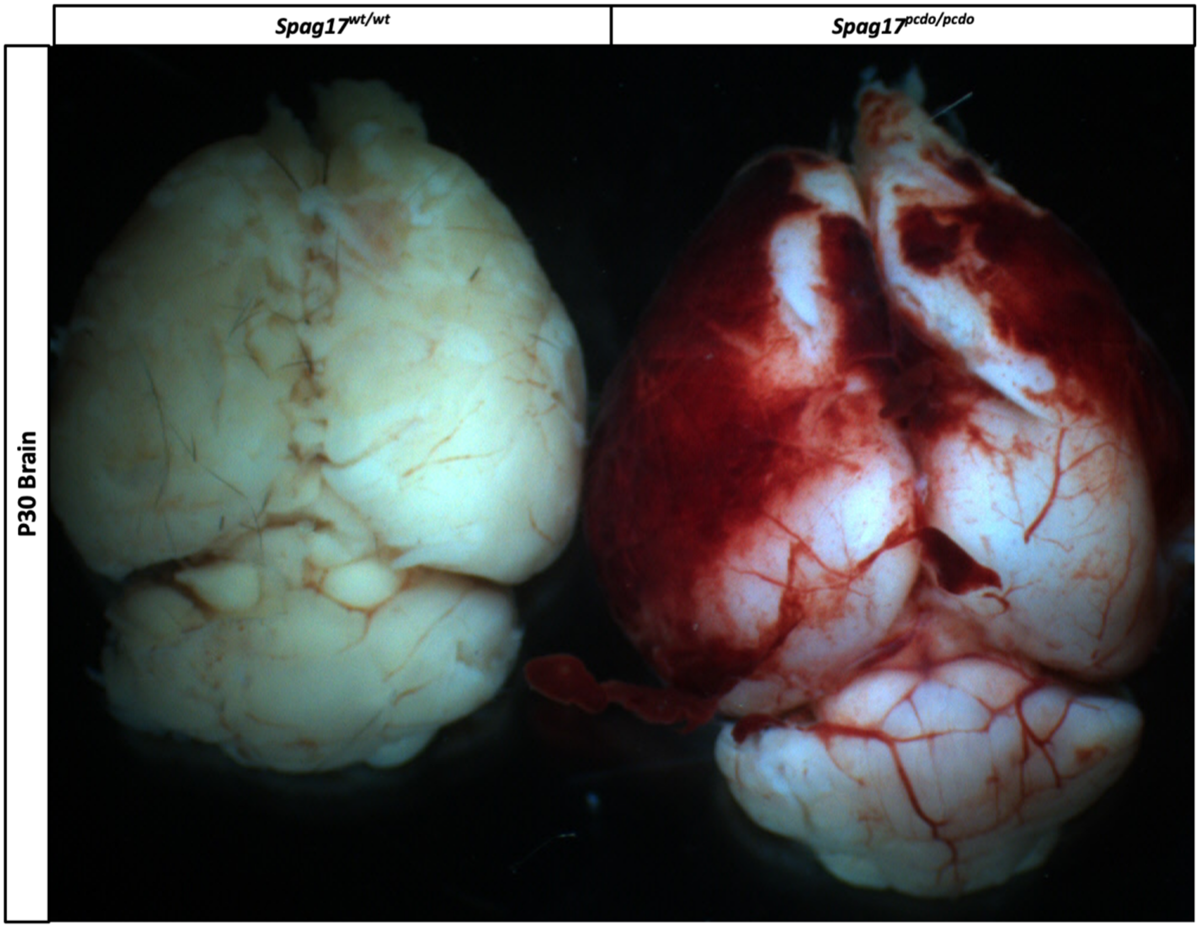

**Supplemental Figure 2:** H&E stained P90 lung sections from animals with the indicated genotypes, showing dilated terminal alveolar ducts (red arrows) in the *Spag17*$^{Pcdo/Pcdo}$ mutant's animals.

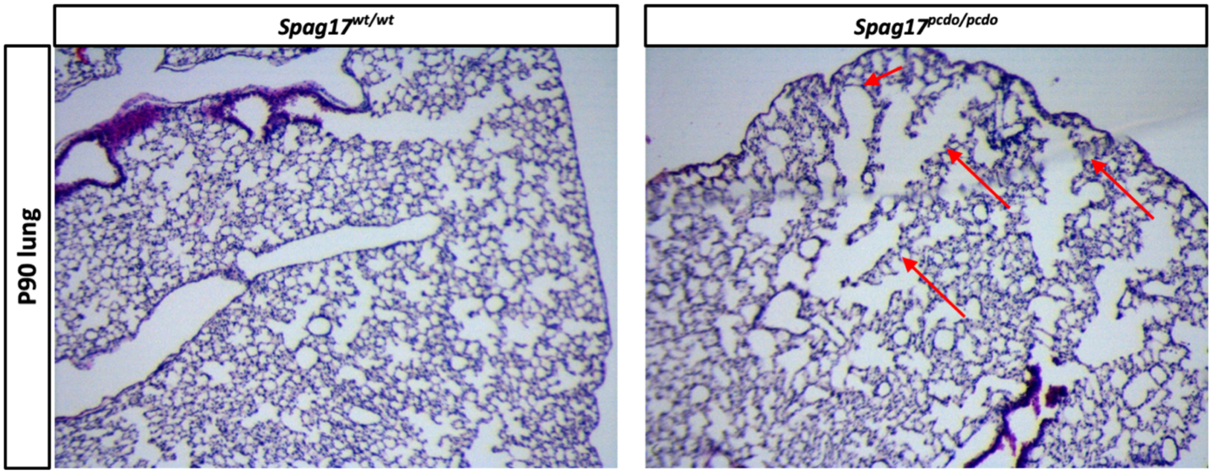

**Supplemental Table 1. Exome sequencing identifies *Spag17* variants in *Pcdo* mutant mice.**

| | Mutant 1 | Mutant 2 | Mutant 3 |
|---|---|---|---|
| Total variants | 118,892 | 126,145 | 114,194 |
| Homozygous | 52,734 | 73,800 | 56,014 |
| Not in dbSNP | 8,665 | 12,779 | 9,874 |
| “high” impact | 10 | 14 | 7 |
| “moderate” impact | 497 | 763 | 638 |
| Common “high” impact | 3: *Plxnc1, Sfi1, Spag17* | | |

**Supplemental Video 1:** High speed video recording of the p4 *Spag17*$^{wt/wt}$ ependymal cilia of the medial wall of the lateral ventricle. 300 frames acquired at a rate of 400 per second.

**Supplemental Video 2:** High speed video recording of the p4 *Spag17*$^{Pcdo/Pcdo}$ ependymal cilia of the medial wall of the lateral ventricle. 300 frames acquired at a rate of 400 frames per second.

**Supplemental Video 3:** Green fluorescent light alternating with DIC white light to generate high speed video microscopy recording that ependymal cilia beating simultaneously with the moving green flourescent beads and the beating cilia along the medial wall of the P4 medial wall of the lateral ventricle of the *Spag17*$^{wt/wt}$ animals. 300 frames captured at a rate of 66 frames per second.

**Supplemental Video 4:** Green fluorescent light alternating with DIC white light to generate high speed video microscopy recording that ependymal cilia beating simultaneously with the moving green flourescent beads and the beating cilia along the medial wall of the P4 medial wall of the lateral ventricle of the *Spag17*$^{Pcdo/Pcdo}$ mutant animals. 300 frames captured at a rate of 66 frames per second.

**Supplemental Video 5:** Green fluorescent light alternating with DIC white light to generate high speed video microscopy recording that ependymal cilia beating simultaneously with the moving green flourescent beads and the beating cilia of the intact aqueduct slice of wild type animals. 300 frames captured at a rate of 66 frames per second.

**Supplemental Video 6:** Green fluorescent light alternating with DIC white light to generate high speed video microscopy recording that ependymal cilia beating simultaneously with the moving green flourescent beads and the beating cilia of the intact aqueduct slice of *Spag17*$^{Pcdo/Pcdo}$ mutant animals. 300 frames captured at a rate of 66 frames per second. The flowing beads seem to move in circle with reduced flow speed.
